# New approaches to species delimitation and population structure of corals: two case studies using ultraconserved elements and exons

**DOI:** 10.1101/2020.04.01.021071

**Authors:** Katie L. Erickson, Alicia Pentico, Andrea M. Quattrini, Catherine S. McFadden

## Abstract

As coral populations decline worldwide in the face of ongoing environmental change, documenting their distribution, diversity and conservation status is now more imperative than ever. Accurate delimitation and identification of species is a critical first step. This task, however, is not trivial as morphological variation and slowly evolving molecular markers confound species identification. New approaches to species delimitation in corals are needed to overcome these challenges. Here, we test whether target enrichment of ultraconserved elements (UCEs) and exons can be used for delimiting species boundaries and population structure within species of corals by focusing on two octocoral genera, *Alcyonium* and *Sinularia*, as exemplary case studies. We designed an updated bait set (29,363 baits) to target-capture 3,040 UCE and exon loci, recovering a mean of 1,910 ± 168 SD per sample with a mean length of 1,055 ± 208 bp. Similar numbers of loci were recovered from *Sinularia* (1,946 ± 227 SD) and *Alcyonium* (1,863 ± 177 SD). Species-level phylogenies were highly supported for both genera. Clustering methods based on filtered SNPs delimited species and populations that are congruent with previous allozyme, DNA barcoding, reproductive and ecological data for *Alcyonium*, and offered further evidence of hybridization among species. For *Sinularia*, results were congruent with those obtained from a previous study using Restriction Site Associated DNA Sequencing. Both case studies demonstrate the utility of target-enrichment of UCEs and exons to address a wide range of evolutionary and taxonomic questions across deep to shallow time scales in corals.

## Introduction

Corals are arguably one of the most ecologically important groups of metazoans on earth. By their ability to produce large colonies by asexual propagation and CaCO_3_ precipitation, they create massive biogenic structures, thereby engineering entire reef-based ecosystems in both shallow and deep waters (Riegl & Dodge, 2019; Roberts, Wheeler, & Freiwald, 2006). These ecological communities are among those most imperiled by climate change, with dire predictions suggesting that they may not survive the next century (Carpenter et al., 2008; Pandolfi, Connolly, Marshall, & Cohen, 2011; Hughes, Bellwood, Connolly, Cornell, & Karlson, 2014). As coral populations and species are in decline, it is important to establish baselines of abundance, distribution, and diversity, all of which rely on accurately delimiting and identifying species.Unfortunately, our current lack of understanding of basic taxonomy and species boundaries in many groups of corals prohibits us from better understanding their roles in ecosystems, and ultimately, from effectively managing their habitats.

Traditionally, morphological traits such as colony growth form and skeletal structure have been used to distinguish corals at all taxonomic levels (Fabricius & Alderslade, 2001; Veron, 2000). At the species level, however, morphological differences can be subtle, and both phenotypic variation and plasticity of growth form often confound assessment of species boundaries (e.g., Forsman, Barshis, Hunter, & Toonen, 2009; Marti-Puig et al., 2014; McFadden et al., 2017; Paz-García, Hellberg, García-de-León, & Balart, 2015). The advent of integrated taxonomic approaches that utilize molecular characters to inform taxonomy has greatly improved our understanding of species boundaries in numerous coral taxa, and led to the recognition of homoplasy in many morphological characters previously considered to be diagnostic of species and higher taxa (Arrigoni et al., 2014; Benayahu, Ofwegen, & McFadden, 2018; Budd & Stolarski, 2009; McFadden et al., 2017). The relative invariance of the mitochondrial genome In corals and other anthozoan cnidarians (e.g., sea anemones) has, however, hindered the application of integrated approaches to species delimitation in this ecologically important group.

As a result of their slow rates of mitochondrial gene evolution coupled with a lack of alternative marker development, both DNA barcoding efforts and studies of phylogeography in most corals lag far behind other groups. The universal COI barcode does not reliably distinguish species within the anthozoan cnidarians (Huang, Meier, Todd, & Chou, 2008; McFadden et al., 2011; Shearer & Coffroth, 2008). No alternative, universally agreed upon single-locus barcode has been proposed or widely used for either the scleractinian corals or related sea anemones, and a multilocus barcode that has been used for the octocorals (*mtMutS* + COI + 28S rDNA) has recognized limitations (McFadden et al., 2011; McFadden, Brown, Brayton, Hunt, & Ofwegen, 2014; Quattrini et al., 2019). Relatively few studies have developed alternative markers such as nuclear gene regions (Ladner & Palumbi, 2012; Prada et al. 2014) or microsatellites (e.g., Baums, Boulay, Polato, & Hellberg, 2012; Gutiérrez-Rodríguez & Lasker, 2004; LeGoff, Pybus, & Rogers, 2004; Quattrini, Baums, Shank, Morrison, & Cordes, 2015) for use in species delimitation or phylogeographic studies. As a result, understanding of population structure, phylogeographic patterns, and the processes driving evolutionary diversification in corals and other anthozoans lags far behind the progress that has been made in other ecologically important marine phyla (Cowman, Parravicini, Kulbicki, & Floeters, 2017; Crandall et al., 2019; Hodge & Bellwood, 2016).

The recent development of genomic sequencing methods holds considerable promise for resolution of species boundaries and phylogeography in corals and other anthozoans. Several studies have now demonstrated that Restriction Site Associated DNA sequencing (RAD-Seq) effectively delimits species when conventionally-used barcodes have failed (Forsman et al., 2017; Herrera & Shank, 2016; Johnston et al., 2017; Pante et al., 2015; Quattrini et al., 2019). However, few studies have yet documented the effectiveness of RAD-Seq for population-level work in anthozoans (Leydet, Grupstra, Coma, Ribes, & Hellberg, 2018; Bracco, Liu, Galaska, Quattrini, & Herrera, 2019). There are also a few inherent disadvantages to RAD-Seq relative to sequence enrichment methods. These include a stark drop in orthologous loci that can be extracted from distantly related taxa (Cairou, Duret, & Charlat, 2013; Viricel, Pante, Dabin, & Simon-Bouhet, 2013), limiting the ability of RAD-Seq to determine deeper phylogenetic relationships, and rendering this method impractical for combining datasets of distantly-related taxa (Harvey, Smith, Glenn, Faircloth, & Brumfield, 2016).

Target capture enrichment of Ultraconserved Elements (UCEs) and exons could be a welcome addition to the methods used for species delimitation and studies of phylogeography and population structure in corals and other anthozoans. UCEs consist of highly conserved regions shared across species, which makes them ideal loci for target-capture enrichment because a single RNA probe set can be applied to a large sample of highly divergent species (Faircloth et al., 2012; McCormack et al., 2012; Quattrini et al., 2018). Recently, Quattrini et al. (2018) demonstrated that UCEs and exons can resolve deeper-level phylogenetic relationships within class Anthozoa, and Quek et al. (2020) applied a similar transcriptome-based target-enrichment approach to resolve genera of scleractinian corals. However, since highly divergent and phylogenetically informative regions flank UCE loci (Faircloth et al., 2012), they have the potential to resolve phylogenetic relationships at shallow evolutionary scales as well. Previous studies have successfully demonstrated that UCE and/or exon data can be used for species delimitation (e.g., Blair et al., 2019; Pie et al., 2019; Zarza et al., 2018) and population genomics (Oswald et al., 2019; Winker, Glenn, & Faircloth, 2018; Zarza et al., 2016) in a wide range of taxa (e.g., spiders, frogs, snakes, birds) across a range of evolutionary time scales. Moreover, the target-capture method has proven successful with highly degraded DNA, such as is often recovered from museum specimens or specimens not preserved specifically for genomics (Derkarabetian et al., 2019; Hawkins et al., 2016; McCormack et al., 2016; Ruane & Austin, 2017). Here, we test the ability of target-capture of UCEs and exons to delimit species and geographically discrete populations of corals, comparing the results directly to previous studies that have applied RAD-Seq, single-locus barcoding and other molecular methods to the same specimens. If this method is in fact successful at delimiting species and populations, then a single comprehensive dataset could be used to simultaneously estimate population structure within a species, discriminate closely related species, and identify phylogenetic relationships across a variety of timescales to help resolve the anthozoan tree of life.

To investigate the utility of UCEs and exons in delimiting coral species and populations, we present two case studies, both of which have been the focus of previous species delimitation efforts using other molecular methods. The first case study is the genus *Alcyonium*, a common soft coral found circumglobally in temperate and cold waters. *Alcyonium* provides an ideal study system because species boundaries among a group of 10 North Atlantic and Mediterranean species have been confirmed previously using a combination of morphological, reproductive and genetic data (McFadden, 1999; McFadden, Donahue, Hadland, & Weston, 2001; McFadden & Hutchinson, 2004). These 10 species (two currently undescribed) belong to three well-defined clades (the *digitatum, coralloides* and *acaule* groups). Pairs of species in each clade nonetheless share identical or nearly identical haplotypes or genotypes at *mtMutS, COI, 28S rDNA* and *ITS*, the markers that have been used most widely for species delimitation and DNA barcoding in octocorals (McFadden et al., 2001, 2011, 2014; McFadden, Sánchez, & France, 2010). Differences in their reproductive biology (McFadden et al., 2001) as well as fixed allelic differences at allozyme loci (McFadden, 1999) suggest that genetically very similar species are nonetheless reproductively isolated. In addition, the population structure of *A. coralloides* spanning the species’ range in the Mediterranean and eastern North Atlantic has been documented previously using allozymes (McFadden, 1999). Therefore, we can evaluate the utility of using UCEs and exons for both population genetic and species delimitation studies of corals using *Alcyonium* as an exemplar.

Our second case study is the genus *Sinularia*, one of the most speciose, widespread and ecologically important soft corals found on shallow-water reefs throughout the Indo-Pacific. As many as 38 morphospecies of *Sinularia* have been found to co-occur on a single reef (Manuputty & Ofwegen, 2007; Ofwegen, 2002), where they contribute to reef formation and biodiversity maintenance. Despite the important roles that they play in coral reef ecosystems, delimitation of closely related *Sinularia* species based on morphological characters and DNA barcodes remains challenging. Two clades in particular, denoted as clades 4 and 5C, each comprise many nominal species whose morphological distinctions are subtle and often confusing (McFadden, Ofwegen, Beckman, Benayahu, & Alderslade, 2009), and DNA barcodes do not always reliably differentiate distinct morphospecies (McFadden et al., 2014; Benayahu et al., 2018). Recently, Quattrini et al. (2019) combined RAD-Seq data with coalescent and clustering methods to delimit *Sinularia* species, also using the results to evaluate the reliability of character-based DNA barcodes in the group. This work has given us the opportunity to use the same individuals as in Quattrini et al. (2019) to determine if species delimitations based on RAD-Seq vs. UCE/exon loci are congruent.

By targeting the same individuals used in these previous studies, we address whether species and population boundaries delimited with target-capture of UCEs and exons are congruent with findings from other methods. Specifically, we re-designed our original anthozoa-v1 bait set (Quattrini et al., 2018) to capture more loci with a higher capture rate in both subclasses of anthozoans, including octocorals (herein) and hexacorals (Cowman et al., 2020). In this study, we then target-captured these loci within two exemplar genera of alcyonacean soft corals and examined phylogenetic and phylogeographic relationships. In addition, we further modified an existing pipeline (Zarza et al., 2016) to call SNPs from UCE and exon loci. Filtered SNPs were then used in Bayesian and multivariate analyses to delimit species within *Alcyonium* and *Sinularia* and to infer population structure within *A. coralloides*.

## Methods

### Case Study 1: Alcyonium

*Alcyonium* specimens were collected between 1989-1996 from locations in the North Atlantic and Mediterranean using SCUBA, with the exception of *A. palmatum* which was obtained from deep water by dredge (McFadden, 1999; McFadden et al., 2001). All samples were preserved in liquid nitrogen immediately after collection and stored in a -80°C freezer. Sixty-eight *Alcyonium* samples that had been included in previous studies were sequenced herein. Samples included at least three specimens from each of the 10 putative species in three distinct clades (*digitatum, coralloides, acaule*) as denoted by *ITS, mtMutS* and/or *COI* barcodes (McFadden et al., 2001, 2011). In addition, a total of 32 specimens of *A. coralloides* that were analyzed with allozymes (McFadden, 1999) were included, including five individuals from each of four locations: two in the Mediterranean, from Marseille, France (*A. coralloides-*M1) and Banyuls-sur-Mer, France (*A. coralloides-*M2); and two in the eastern North Atlantic, from Iles Chausey, France (*A. coralloides-*CH) and Ilhas do Martinhal, Sagres, Portugal (*A. coralloides-*SG). *Alcyonium haddoni*, collected from Argentina, was also included as it had been sequenced in Quattrini et al. (2018).

### Case Study 2: Sinularia

*Sinularia* from clades 4 and 5C were collected using SCUBA at 13 sites in the shallow fore-reef zone (3-21m) at Dongsha Atoll Marine National Park (Taiwan) in 2011, 2013, and 2015 (Benayahu et al., 2018; Quattrini et al., 2019). All specimens were preserved in 70% EtOH, with small tissue subsamples preserved in 95% EtOH for DNA (Quattrini et al., 2019). Benayahu et al. (2018) identified these individuals to species using morphological characters and DNA barcodes. Subsequently, Quattrini et al. (2019) used RAD-Seq from 95 individuals to confirm species boundaries among morphospecies. A subset of 13 individuals of four morphospecies in clade 4 and 31 individuals of nine morphospecies in clade 5C were selected for sequencing using target-capture methods.

### RNA Bait Design

To have higher specificity for and target more loci in specific taxonomic groups, we re-designed a bait set (anthozoa-v1, Quattrini et al., 2018) that was produced to target-capture 720 UCE and 1,071 exon loci from across the sub-class Anthozoa. Target-capture results with the original bait set from 235 anthozoans (unpubl. data) were screened to keep baits that either performed well across all anthozoans or within particular groups more specifically. Thus, we retained 12,320 baits targeting 1,580 loci (667 UCE and 913 exon loci) from the anthozoa-v1 bait set. Using the program Phyluce (Faircloth, 2016) and following methods in Quattrini et al., (2018), we added more octocoral-specific baits to 1) improve target-capture performance of the anthozoa-v1 loci in octocorals and 2) target additional loci, which were not included in the anthozoa-v1 bait set. A similar procedure was followed for hexacorals, as reported elsewhere (Cowman et al., 2020). Methods are outlined in Suppl. File 1, but more details can also be found in both Quattrini et al. (2018) and the PHYLUCE documentation (https://phyluce.readthedocs.io/en/latest/tutorial-four.html, Faircloth, 2016).

Following bait design, the exon and UCE bait sets were concatenated and then screened to remove duplicate baits (≥50% identical over >50% of their length), resulting in a final non-duplicated octocoral-v2 bait set of 29,363 baits targeting 3,040 loci (1,340 UCE and 1,700 exon loci). Baits designed against transcriptomes have “design: octotrans” in the .fasta headers. It is possible that both “UCE” loci and “exon” loci contain both conserved coding and non-coding regions, but we retain the distinction here based on how the baits were designed and following naming conventions in prior studies (Branstetter, Longino, Ward, & Faircloth, 2017; Starrett et al., 2017; Quattrini et al., 2018). Baits were synthesized by Arbor BioSciences (Ann Arbor, MI).

### DNA extractions, library preps, and sequencing

DNA was extracted using the Qiagen DNeasy Blood & Tissue kit or a modified CTAB extraction protocol (McFadden et al., 2001). Approximately 250 ng of DNA from each sample was then sheared to an average size of 400-800 bp (Q800R QSonica Inc. Sonicator). Libraries were prepared for sequencing using the KAPA HyperPrepKit (Kapa Biosystems) with dual-indexed iTru adapters following Quattrini et al. (2018). For 66 *Alcyonium* samples and 38 *Sinularia* samples, the MyBaits protocol v IV (Arbor BioSciences, Ann Arbor, MI) was used to target and enrich UCEs and exons in pools of eight individuals using the newly designed bait set. For three *Alcyonium* and six *Sinularia*, loci were captured with the anthozoa-v1 bait set as described in Quattrini et al. (2108). Library size was assessed using a Bioanalyzer (UC Riverside) and then sent to Oklahoma Medical Research Foundation (Oklahoma City, OK) for sequencing. Samples target-enriched with the anthozoa-v1 bait set were sequenced with other specimens on an Illumina HiSeq 3000 (120 samples multiplexed) and samples target-enriched with the newly designed octocoral-v2 bait set were sequenced on a NovaSeq (214 samples multiplexed). More methodological details can be found in Quattrini et al. (2018).

### Bioinformatic processing, assembly, and alignment of UCEs

The PHYLUCE pipeline (Faircloth, 2016) was used to perform quality control on the demultiplexed reads, assemble loci, and align loci for phylogenetic analyses. Reads were cleaned using illumiprocessor (Faircloth, 2013) and Trimmomatic v 0.35 (Bolger, Lohse, & Usadel, 2014). Contigs were assembled using the program SPAdes v 3.1 (Bankevich et al., 2012; with the --careful and --cov-cutoff 2 parameters). The *phyluce_assembly_match_contigs_to_probes* command was used to match probes to contigs to identify loci with a minimum coverage of 70% and a minimum identity of 70%. Loci were then extracted using *phyluce_assembly_get_fastas_from_match_counts* and aligned with MAFFT v7.130b (Katoh et al., 2002). Edges of the aligned loci were then trimmed using *phyluce_align_seqcap_align*. Coverage of each locus was estimated using *phyluce_assembly_get_trinity_coverage* and *phyluce_assembly_get_trinity_coverage_for_uce_loci*. Finally, a data matrix including all loci concatenated with at least 75% of taxa present was generated using the program *phyluce_align_get_only_loci_with_min_taxa*.

### Phylogenetics

Maximum likelihood inference was run on concatenated alignments of the 75% complete UCE and exon data matrix to obtain a phylogenetic tree. Phylogenies were constructed using the GTRGAMMA model by searching for the maximum-likelihood (ML) tree. The program RAxML was used to perform 100 bootstrap searches and 20 ML searches using rapid bootstrapping (Stamatakis, 2014). A phylogeny was constructed for the *Alcyonium* samples along with three *Sinularia* spp. as outgroups. RAD-Seq data from *Sinularia* as described in Quattrini et al. (2019) were subset to include only the individuals used in target-capture analysis. ML was performed on both RAD-Seq and UCE/exon datasets using the above methods on clades 4 and 5C to determine the congruence between constructed phylogenies. Clade 4 was rooted to *S. humilis* while a midpoint root was used for Clade 5C as in Quattrini et al. (2019).

### SNP calling

In order to conduct population level analyses, SNPs were extracted from each clade. In each taxon set, the individual with the largest number of UCE/exon contigs as identified by *phyluce_assembly_get_match_counts* was chosen as a reference individual for SNP calling. The programs *phyluce_assembly_get_match_counts* and *phyluce_assembly_get_fastas_from_match_count* were re-run on the reference individual to create a fasta of UCE and exon contigs found just in the reference. Reference loci were indexed using bwa-version 0.7.7 (Li & Derbin, 2009). BAM files for each individual in the analysis were created by mapping their reads to the reference using BWA-MEM (Li, 2013), sorting the reads using SAMtools and removing duplicates using Picard v 1.106-0 (http://picard.sourceforge.net). GATK v3.4 (McKenna et al., 2010) was then used to realign BAMs around indels, call variants, and filter variants based on vcf tools (Danecek et al., 2011). For each clade, SNPs were called at loci for which at least 75% of taxa were represented and for which at least 10 reads mapped to each locus. Only individuals target-captured with the new octocoral-v2 bait set were included in SNP calling analyses to maximize number of SNPs called. SNPs were filtered to one SNP per locus for loci ≤ 1,000 bp, or to one per 1,000 bp if loci were ≥ 2,000 bp. All code for SNP calling can be found in Supplemental File 2.

### Species delimitation

Two methods were used to delimit species: Discriminant Analysis of Principal Components (DAPC) and STRUCTURE. Both of these programs used the filtered SNPs to identify clusters of genetically-related individuals for each clade. DAPC (Jombart et al., 2010) clusters species based on their genetic similarity and can provide estimates for how many populations (*K*) to assume. DAPC was conducted using the adegenet package (Jombart, 2008) in R (R Core Team, 2019). First the program *find*.*clusters* found the optimal number of clusters needed to minimize the Bayesian Information Criterion (BIC), and the function *optim*.*a*.*score* was used to determine how many Principal Component (PC) axes and Discriminant Functions (DFs) needed to be retained. The DAPC results were displayed as scatterplots.

STRUCTURE (Pritchard, Stephens, & Donnelly, 2000) uses a Bayesian clustering approach to probabilistically infer population structure given *K*. STRUCTURE was run in parallel using StrAutoParallel (Chhatre & Emerson, 2017, generations=1M, burnin=250K). The maximum value for *K* was chosen to be higher than the optimal number of clusters designated by DAPC. Five runs were completed at each value of *K*. The five runs at the DAPC-selected *K* value were combined and results were visualized using *starmie* (https://github.com/sa-lee/starmie) in R (R Core Team, 2019).

## Results

### Sequencing and Locus Summary

The total number of reads obtained from the HiSeq ranged from 2,680,770 to 9,150,274 and the NovaSeq from 696 to 26,147,550 total reads per sample. We removed two *Alcyonium* samples that failed sequencing, and then quality and adapter trimmed raw reads for the remaining samples, resulting in a mean of 8,021,057 ± 83,566 SD PE trimmed reads per sample (Supplemental Table 1). Trimmed reads were assembled into a mean of 83,566 ± 57,749 SD contigs per sample (range: 22,813 to 285,695) with a mean length of 400 ± 32 bp (Supplemental Table 1). Read coverage per contig ranged from 0.4 to 5.5X with the original bait set and HiSeq sequencing and 0.1 to 9.6X with the new bait set and NovaSeq sequencing.

**Table 1.** Number of SNPs recovered for each clade.

|  | # Unfiltered SNPs | # Filtered SNPs |
| --- | --- | --- |
| *Alcyonium* |  |  |
| *digitatum* clade | 30,384 | 1,974 |
| *coralloides* clade | 39,302 | 1,980 |
| *acaule* clade | 37,569 | 2,114 |
| *coralloides* populations | 4,775 | 152 |
| *Sinularia* |  |  |
| Clade 4 | 65,261 | 2,030 |
| Clade 5C | 71,400 | 2,742 |

With the new bait set and NovaSeq sequencing, a total of 2,977 loci (out of 3,040 targeted loci) were recovered from the assembled contigs (Supplemental Table 1). The mean number of loci recovered with this new bait set was 1,910 ± 168 SD per sample (range: 1,019 to 2,173) with a mean length of 1,055 ± 208 bp (range:115 to 1,803 bp). This method produced an average of 2X more loci that were 200 bp longer as compared to target capture with the anthozoa-v1 bait set (994 ± 194 SD loci recovered at 886 ± 191 bp) and sequencing with the HiSeq. We recovered similar numbers of loci from *Sinularia* (1,991 ± 114 SD) as compared with *Alcyonium* (1,863 ± 177 SD). Read coverage per UCE contig ranged from 1 to 12.8X with the original bait set and HiSeq and 1.2 to 135X with the new bait set and NovaSeq.

### Case Study 1: Alcyonium

The 75% taxon occupancy matrix included a concatenated alignment of 1,264 UCE and exon loci with an alignment length of 1,280,803 bp. ML inference on this dataset produced a highly supported phylogeny (most nodes >95% b.s.), except for the sister relationship between the *A. digitatum* clade and *A. haddoni* (15% b.s. support) (Fig. 1). In the *digitatum* clade, each putative species (*A*. sp. A, *A. digitatum, A. siderium)* was monophyletic. In the *coralloides* clade, most of the putative species were monophyletic, except that one *A*. sp. M2 was sister to the remaining species in the *coralloides* clade. In the *acaule* clade, *A. acaule* was monophyletic but *A. glomeratum and A. palmatum* were nested together.

**Figure 1.**
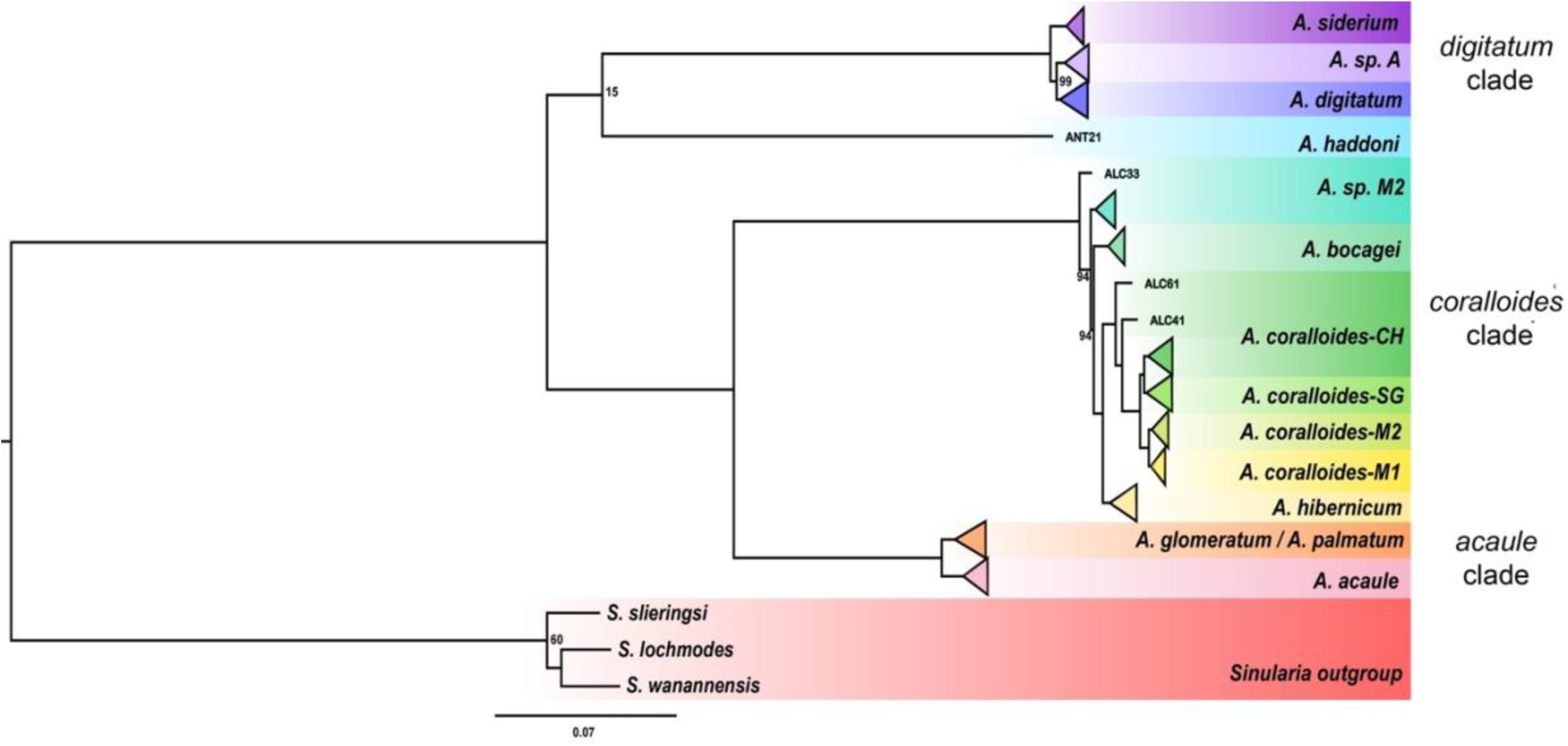
Maximum likelihood phylogeny of *Alcyonium* based on a 75% complete matrix containing 1,264 UCE loci (1,280,803 bp). All unlabeled nodes have 100% bootstrap support (100 b.s. replicates). The tree was rooted to *Sinularia* outgroups.

Both DAPC and STRUCTURE analyses based on filtered SNPs supported the phylogenetic results for each group (Fig. 2). For the *digitatum* clade, DAPC (1,974 filtered SNPs) indicated *K*=3 clusters matching the three putative species; however, STRUCTURE (*K*=3) indicated that a few *A*. sp. A individuals were admixed with *A. digitatum* (Fig. 2A-B). For the *coralloides* clade, DAPC (1,980 filtered SNPs) suggested *K*=4 clusters, with Mediterranean *A. coralloides*, north Atlantic *A. coralloides, A. bocagei*, and *A. hibernicum/*sp. M2 forming distinct clusters (Fig. 2C-D). STRUCTURE at *K*=4 also revealed that *A. hibernicum* was admixed between north Atlantic *A. coralloides* and *A*. sp. M2. For the *acaule* clade, DAPC (2,114 filtered SNPs) indicated *K*=3 clusters corresponding to the three putative species; however, STRUCTURE at *K*=3 indicated that all *A. glomeratum* individuals were admixed with *A. palmatum* (Fig. 2E-F).

**Figure 2.**
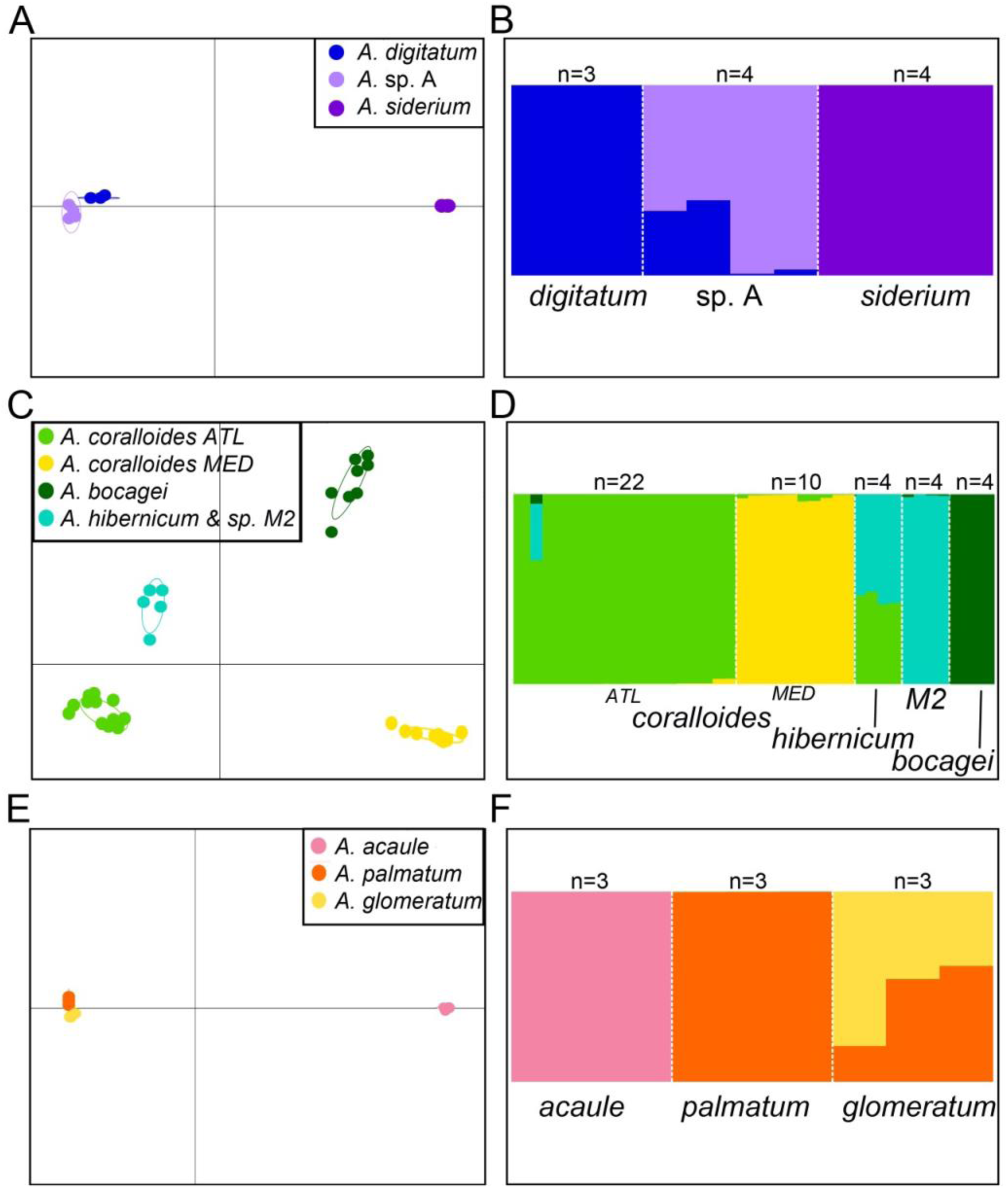
Probability of membership graph from STRUCTURE analyses and Discriminant Analysis of Principal Components (DAPC) plot for (A-B) *Alcyonium digitatum* clade (1,974 SNPs, *K*=3), (C-D) *Alcyonium coralloides* clade (1,980 SNPs, *K*=4), and (E-F) *Alcyonium acaule* clade (2,114 SNPs, *K=3)*.

We also examined whether this method could be used to discern populations of *A. coralloides* in the Mediterranean Sea from those in the eastern North Atlantic (Fig. 3A). Although 4,775 unfiltered SNPs were recovered from individuals used in analyses (Table 1), only 152 filtered SNPs were obtained. Nevertheless, the two populations in the Atlantic (*CH, SG*) were distinct from one another and were clearly separated from those in the Mediterranean (*M1, M2*). In addition, the Mediterranean populations were differentiated from one another, although a few individuals were admixed (Fig. 3B, C).

**Figure 3.**
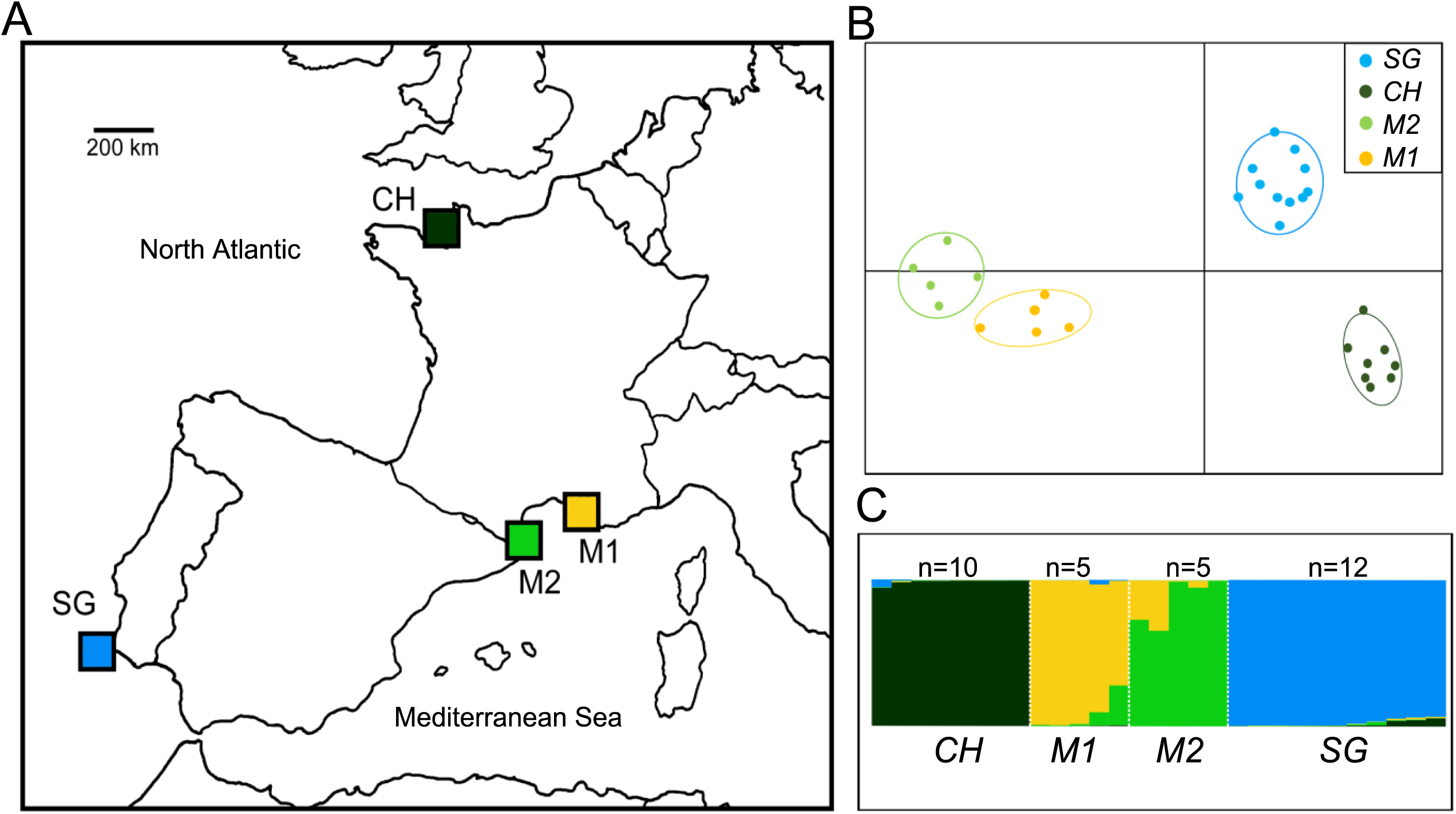
(A) Map of collection locations for *Alcyonium coralloides*, (B) Probability of membership graph from STRUCTURE analyses and (C) Discriminant Analysis of Principal Components (DAPC) plot for *A. coralloides* (152 SNPs, *K*=4) populations (M1: Marseille, France, M2: Banyuls-sur-Mer, France, CH: Iles Chausey, France, SG: Ilhas do Martinhal, Sagres, Portugal).

### Case Study 2: Sinularia

For *Sinularia* clade 4, the 75% taxon occupancy matrices included concatenated alignments of 2,231,641 bp (1,939 loci) and 549,862 bp (6,343 loci) for target-capture and RAD-Seq, respectively. The clade 4 tree topologies for the target-capture and RAD-Seq datasets were identical, with highly supported (> 95% b.s.) monophyletic clades for five putative species (Fig. 4). The concatenated alignments for *Sinularia* clade 5C were 1,549,273 bp (1,240 loci) and 701,433 bp (8,060 loci) for target-capture and RAD-Seq datasets, respectively. Clade 5C tree topologies for both datasets were also highly supported with monophyletic clades for seven putative species; however, *S. penghuensis* and *S. slieringsi* formed a clade and *S. acuta* and *S. abrupta* were nested together (Fig. 5). There was also a notable difference between topologies: the clade containing *S. acuta* and *S. abrupta* was sister to *S. slieringsi/penghuensis* in the RAD-Seq phylogeny, but sister to the other five species in the phylogeny produced with target-capture data. Of note, these phylogenies were rooted at the mid-point. In addition, the relationships among individuals within the clade of *S. lochmodes* differed between datasets; however, these relationships were poorly supported in both RAD-Seq and target-capture phylogenies.

**Figure 4.**
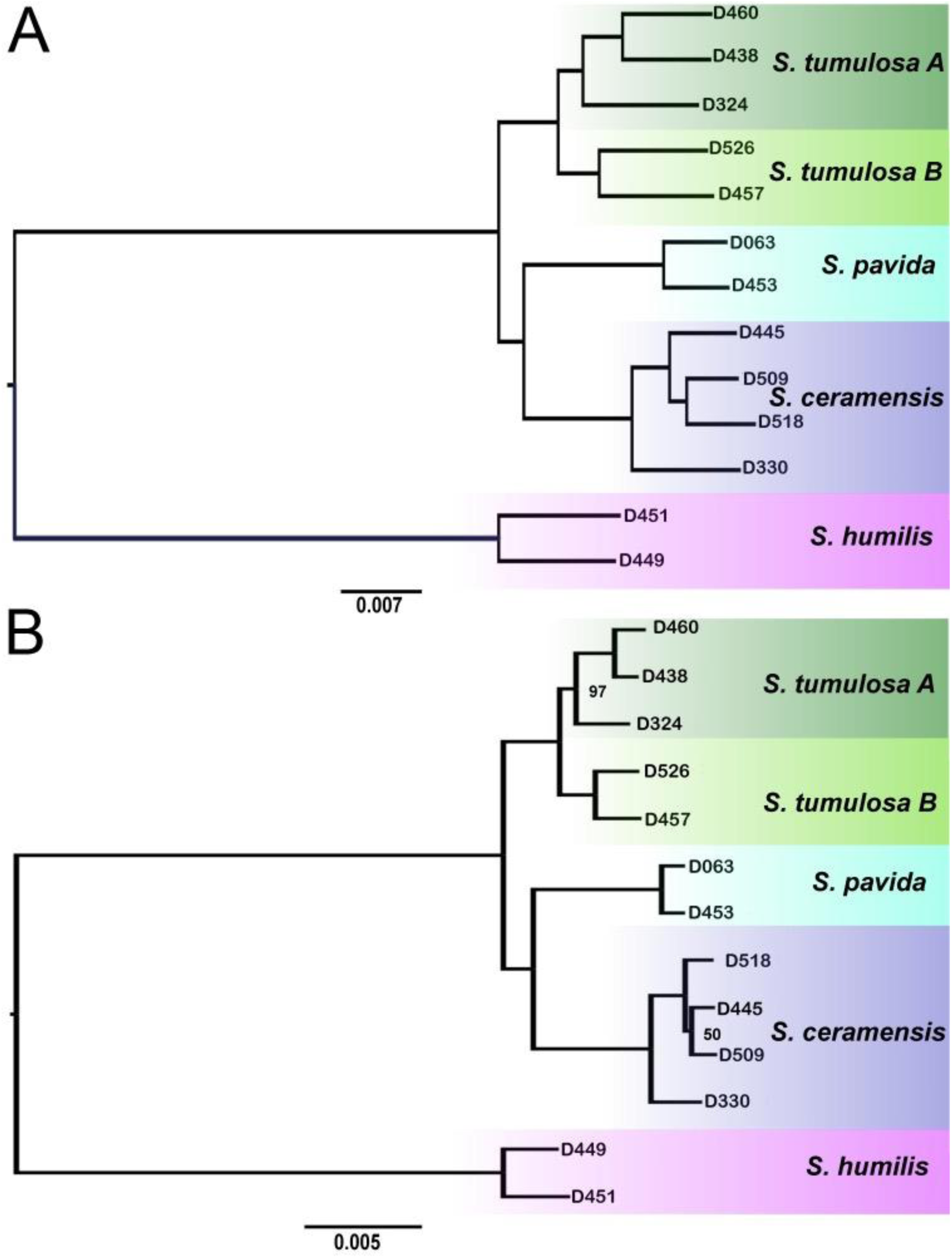
Maximum likelihood phylogenies of *Sinularia* clade 4 constructed from (A) UCE loci (1,939 loci, 2,231,641 bp) and (B) RAD-Seq loci (6,343 loci, 549,862 bp) based on 75% incomplete matrices. All unlabeled nodes have 100% bootstrap support (100 b.s. replicates). The phylogeny was rooted to *S. humilis*.

**Figure 5.**
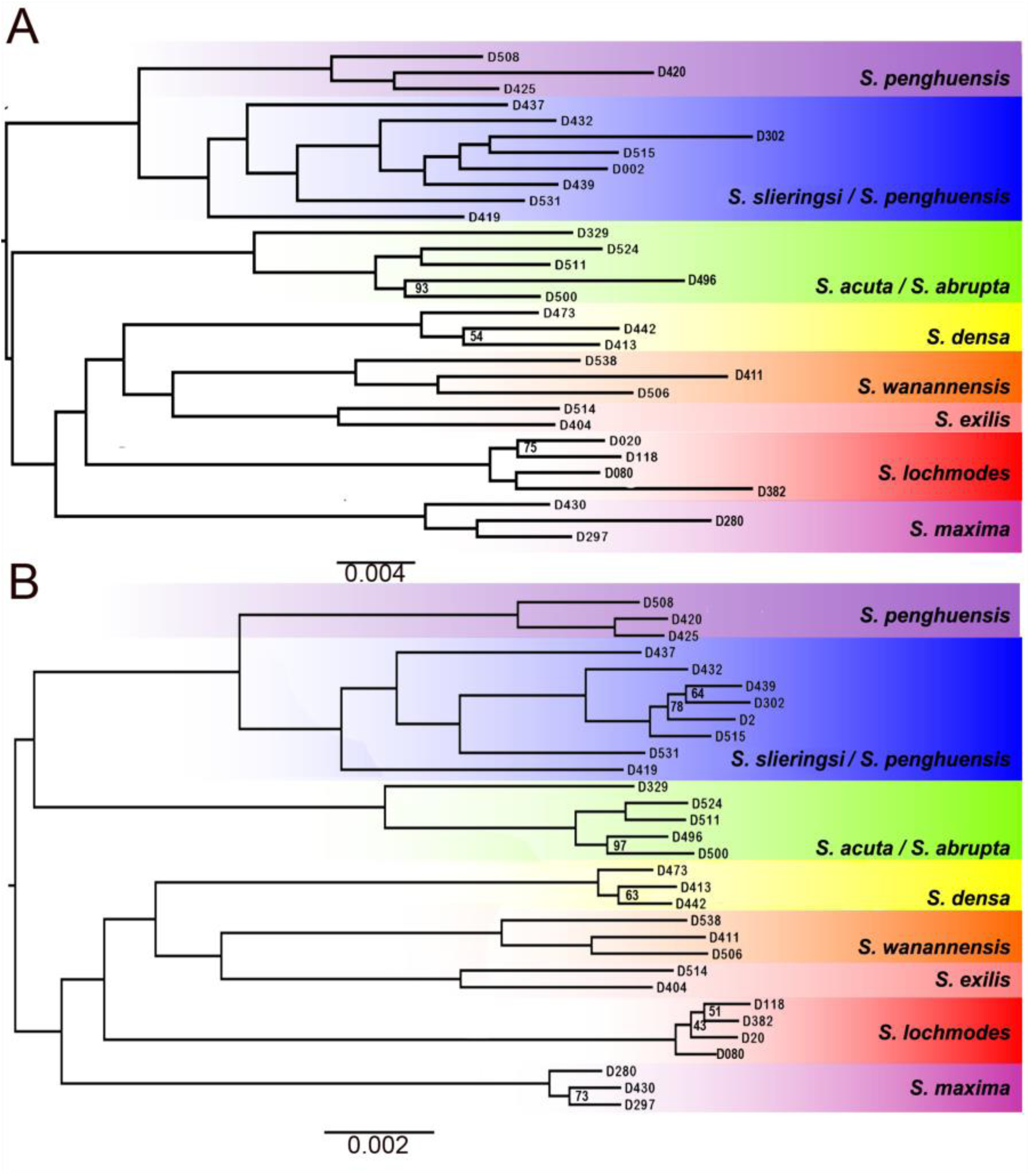
Maximum likelihood phylogenies of *Sinularia* clade 5C constructed from (A) UCE loci (1,240 loci, 1,549,273 bp) and (B) RAD-Seq loci (8,060 loci, 701,433 bp) based on 75% incomplete matrices. All unlabeled nodes have 100% bootstrap support (100 b.s. replicates). A midpoint rooting was used.

We also extracted SNPs from the UCE and exon loci for both *Sinularia* clades. For clade 4, 2,030 filtered SNPs were used in the DAPC and STRUCTURE analyses. DAPC indicated that *Sinularia* clade 4 contains *K*=5 clusters: *S. ceramensis, S. humilis, S. pavida, S. tumulosa A*, and *S. tumulosa B* (Fig. 6A). However, one *S. tumulosa* A individual grouped with *S. tumulosa* B and one *S. ceramensis* individual grouped with *S. humilis*. Plotting *K=5* clusters in the STRUCTURE analysis revealed four species that were well separated: *S. humilis, S. pavida, S. tumulosa* A, and *S. tumulosa* B, except for the one *S. tumulosa* A that was admixed with *S. tumulosa* B. Although the DAPC plot separated most *S. ceramensis* from the remaining species, the STRUCTURE plot suggested all *S. ceramensis* individuals were admixed with *S. humilis*.

**Figure 6.**
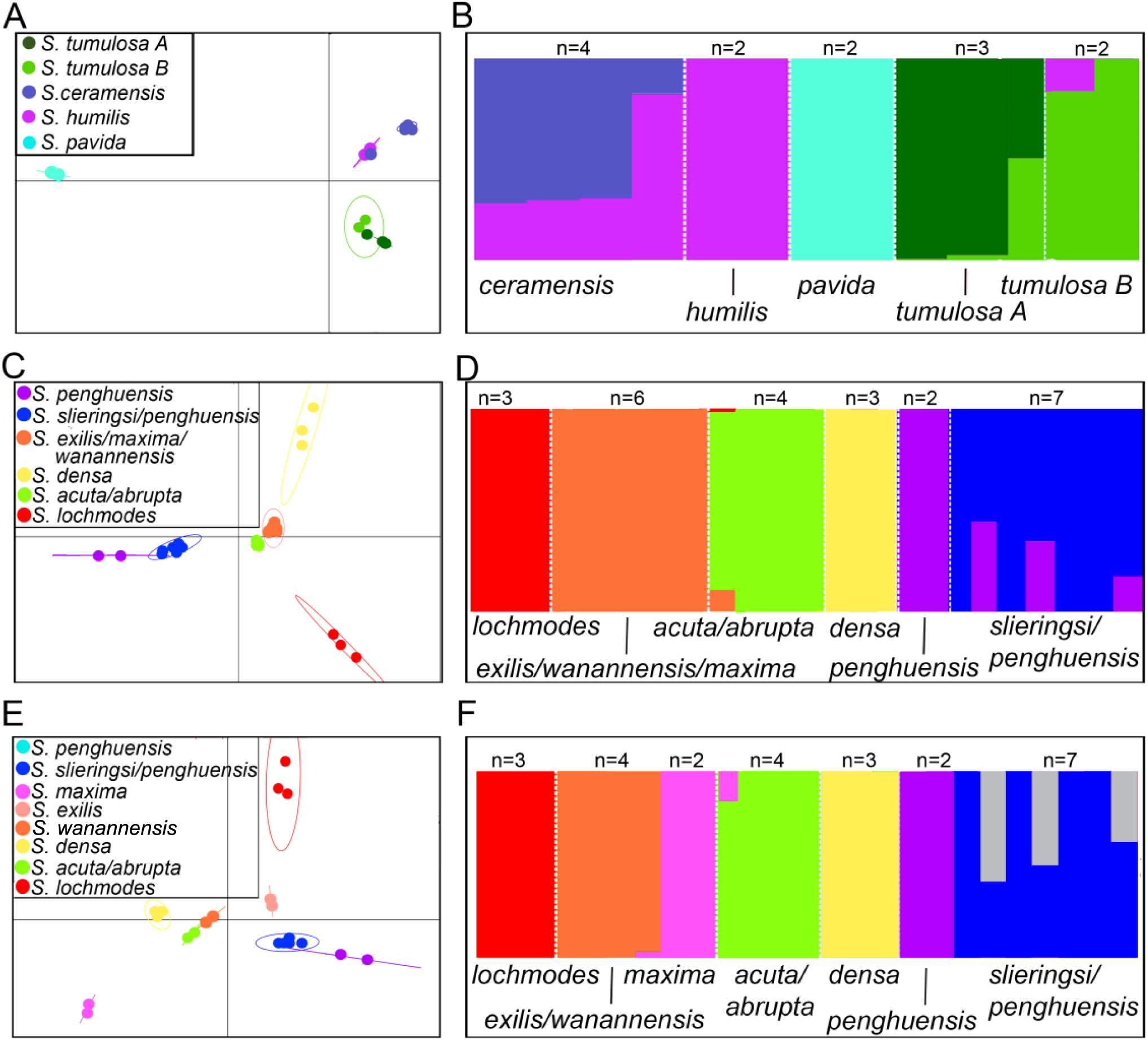
Probability of membership graph from STRUCTURE analyses and Discriminant Analysis of Principal Components (DAPC) plot for (A-B) *Sinularia* clade 4 (2,030 SNPs, *K*=5) and (C-D) *Sinularia* clade 5C (2,742 SNPs, *K*=6), and (E-F) *Sinularia* clade 5C (2,742 SNPs, *K*=8).

The DAPC analysis indicated *K*=6 clusters for *Sinularia* Clade 5C based on 2,742 SNPs (Fig. 6C). Five of the six clusters corresponded to the prior group designations from RAD-Seq analyses (Quattrini et al., 2019): *S. slieringsi/penghuensis, S. penghuensis, S. densa, S. abrupta/acuta*, and *S. lochmodes*. The sixth group consisted of three putative species: *S. exilis, S. maxima*, and *S. wanannensis*. The STRUCTURE plot also indicated clear separation of the six groups, with some admixture observed in *S. slieringsi/penghuensis* individuals with *S. penghuensis* (Fig. 6D). We also plotted clusters at *K*=8 in both DAPC and STRUCTURE analyses based on results of RAD-Seq data (Quattrini et al., 2019). The DAPC plot separated each putative species, but the STRUCTURE analysis indicated that *S. wanannensis* and *exilis* clustered together, but were distinct from *S. maxima* (Fig. 6E,F).

## Discussion

In the present study, we designed a bait set to improve target-capture performance of the anthozoa-v1 bait set (Quattrini et al., 2019) and target additional UCE and exon loci in octocorals (octocoral-v2, 29,363 baits targeting 3,040 loci). Testing this bait set in two well-studied octocoral genera demonstrated the utility of target-capture genomics to delimit both species and populations of octocorals. To date, only three studies have used genomic approaches to delimit among closely-related octocorals, and all three used RAD-Seq data (Herrera & Shank, 2016; Pante et al., 2015; Quattrini et al., 2019), which precludes combining or including other taxonomic groups in these datasets. Here, we demonstrated that phylogenies constructed with RAD-Seq or target-capture UCE/exon data were congruent and contained similar levels of resolution. This target-capture approach was also able to confirm species boundaries, known from reproductive and allozyme data, that were only weakly supported by the widely used DNA barcodes for octocorals (e.g., McFadden et al., 2010). Our enhanced octocoral bait set combined with the companion hexacoral bait set (Cowman et al., 2020) are now available to help resolve the anthozoan tree of life across deep to shallow evolutionary time scales.

### Case Study 1: Alcyonium

*Alcyonium* was one of the first coral genera for which molecular evidence was used to delimit species (McFadden, 1999; McFadden et al., 2001), and its northern hemisphere representatives have continued to serve as test cases for the development of markers for DNA barcoding (McFadden et al., 2011, 2014). Previous studies have used allozymes and various single-locus barcoding markers to: (1) identify two new, cryptic species within the genus (McFadden, 1999; McFadden et al. 2001); (2) support species boundaries among eight of the 10 putative species belonging to three distinct clades (McFadden, 1999; McFadden et al. 2001, 2011, 2014); and (3) suggest that hybridization may have occurred in the past or be ongoing among species in two of those clades (McFadden & Hutchinson, 2004; McFadden, Rettig, & Beckman, 2005). The results of species delimitation using evidence from sequence capture of UCEs and exons provide strong support for each of these hypotheses. Moreover, they strengthen and corroborate formerly weak evidence suggesting: (4) a species boundary between north Atlantic and Mediterranean populations of *A. coralloides* (McFadden, 1999), and (5) no genetic differentiation between *A. palmatum* and *A. glomeratum* (McFadden et al. 2001, 2011, 2014).

The four species comprising the *A. coralloides* clade formerly were synonymized under that name (Groot & Weinberg, 1982), but fixed allelic differences at four to six allozyme loci between sympatrically occurring morphotypes support the distinction of *A. hibernicum, A. bocagei* and *A*. sp. M2 (an undescribed species from the Mediterranean) from *A. coralloides sensu stricto* (McFadden, 1999). Molecular evidence for a species boundary between distinctly different north Atlantic and Mediterranean morphotypes of *A. coralloides*, has, however, been equivocal (McFadden, 1999; McFadden et al., 2011). Our phylogenetic results indicate monophyly for each of these species, with the exception of one individual of *A*. sp. M2 that was recovered sister to the rest of the *coralloides* clade. STRUCTURE and DAPC analyses also support species boundaries between *A*. sp. M2, *A. bocagei*, and the north Atlantic and Mediterranean morphotypes of *A. coralloides*. However, DAPC analysis grouped *A. hibernicum* with *A*. sp. M2, and STRUCTURE results indicated that all (n=4) *A. hibernicum* individuals were admixed between *A*. sp. M2 and *A. coralloides*. Based on observations of shared *ITS* polymorphisms, McFadden & Hutchinson (2004) proposed that *A. hibernicum*, a putatively asexual species (Hartnoll, 1977), arose as the product of hybridization between *A. coralloides* and *A*. sp. M2. They could not, however, conclusively rule out the possibility that these three species might share *ITS* sequences due to recent ancestry and incomplete lineage sorting. STRUCTURE analysis demonstrated that *A. hibernicum* was equally admixed between *A*. sp. M2 and the Atlantic morphotype of *A. coralloides*, supporting the hypothesis that *A. hibernicum* is a hybrid species that arose from a cross between those two species. Because the present-day distributions of the two parental species are allopatric, this event likely happened at some time in the past, perhaps prior to *A*. sp. M2 (re)colonizing the Mediterranean subsequent to the Messinian crisis (Duggen et al., 2003).

Whereas the hybridization event that gave rise to the hybrid species, *A. hibernicum*, likely occurred long ago, it has also been proposed that hybridization may be ongoing within the *digitatum* clade, between *A. digitatum* and *A*. sp. A, an undescribed species from the British Isles (McFadden et al., 2001; McFadden et al., 2005). Where these two sympatric species co-occur in close proximity, individuals that are intermediate in morphology are occasionally found. Genetic evidence from a single allozyme locus and from shared polymorphisms in *ITS* are consistent with the conclusion that these intermediate individuals are F1 hybrids (McFadden et al., 2005), but the extreme genetic similarity of the two parent species at all other loci that have been examined to date made it difficult to establish that fact with certainty. Although only two putative hybrid individuals with intermediate morphologies were sequenced in this study, STRUCTURE analyses indicated that both of those individuals appear to be admixed equally with *A. digitatum* and *A*. sp. A. Sequencing of additional hybrid and parental individuals will be necessary to confirm the status of the hybrids as F1s, and to support the hypothesis that they propagate asexually (McFadden et al., 2005).

Within the *acaule* clade, *A. palmatum* and *A. glomeratum* are distinct from one another morphologically, geographically and ecologically, yet share identical haplotypes or genotypes at *ITS, 28S rDNA, mtMutS* and *COI* (McFadden et al., 2001, 2011, 2014). *A. palmatum* is restricted to deep-water soft sediment habitats in the Mediterranean and north Atlantic, while *A. glomeratum* occurs on rocky substrates in shallow water in the north Atlantic only (Manuel, 1981). In addition to these ecological differences, *A. palmatum* and *A. glomeratum* also differ in colony growth form, with *A. glomeratum* more closely resembling *A. acaule*, an ecologically similar species found on hard substrate in the Mediterranean (Weinberg, 1977). As in previous studies (McFadden et al. 2001, 2011), our phylogenetic and species delimitation analyses indicated that *A. acaule* was distinct from *A. glomeratum* and *A. palmatum*, but the evidence that *A. glomeratum* and *A. palmatum* are distinct species is less clear. Although they formed separate clusters in DAPC analysis, neither species was monophyletic and STRUCTURE analysis indicated admixture. Recent sequence data from an Atlantic population of *A. palmatum* support the hypothesis of ongoing gene flow between these two species (unpubl. data), but more extensive sampling will be required to confirm that possibility.

The target-capture approach also enabled us to differentiate among populations of *A. coralloides*, even with relatively few SNPs (152). The STRUCTURE and DAPC analyses revealed differentiation between the two north Atlantic populations (∼1500 km apart) and between the two Mediterranean populations (∼200 km apart); only a few individuals in the latter populations were admixed. Furthermore, analyses clearly differentiated the Mediterranean and Atlantic populations of *A. coralloides* and indicated no admixture, suggesting that they may indeed be different species. Results correspond to previous analyses using allozymes (McFadden, 1999), and thus show potential in helping to discern populations in other species of corals.

### Case Study 2: Sinularia

Phylogenies of *Sinularia* clade 4 and 5C constructed with target-capture UCE/exon data were congruent with those based on RAD-Seq data (Fig. 4-5 in Quattrini et al., 2019). Both analyses used 75% taxon-occupancy matrices, and recovered highly resolved trees, with the majority of nodes having 95-100% bootstrap support. Only a few nodes in each analysis had <95% b.s. support, although there was a trend for more highly supported nodes in the phylogeny constructed with UCE/exon data. Overall, the target-capture data consisted of fewer loci (∼2K loci) compared to RAD-Seq data (∼6-8k loci), but average locus size (1K bp) and alignments were longer (∼1-2M bp) in UCE/exon data as compared to RAD-Seq data (∼87 bp locus size, 500-700K bp alignment size). One observed difference between the clade 5C phylogenies was the placement of the *S. acuta/abrupta* clade, with this clade sister to *S. penghuensis/slieringsi* in the RAD-Seq phylogeny and sister to the remaining species in the UCE/exon phylogeny. This difference might be attributed to the fact that we used midpoint rooting in the analyses herein. However, using midpoint rooting on an expanded dataset, Quattrini et al. (2019) also found incongruence in the placement of this clade depending on method used (e.g., SNAPP using SNPs or ML using concatenated data) and also indicated that some members of this clade were hybrids. Regardless of the unresolved placement of *S. acuta/abrupta*, topologies based on UCEs/exons were highly congruent with those produced from RAD-Seq data, further supporting that the method (RAD-Seq vs. UCE target capture) does not influence phylogenetic results (Harvey et al., 2016; Manthey et al., 2016).

Our results also suggest that the effectiveness of target enriching UCEs/exons for species delimitation is similar to the more frequently used RAD-Seq approach. DAPC and STRUCTURE results were generally congruent with those from RAD-Seq (Quattrini et al. (2019), although there were a few differences. Within clade 4, three clusters present in the DAPC analysis corresponded to groups designated in Quattrini et al. (2019) by DAPC and STRUCTURE analyses: S. *humilis, S. pavida*, and *S. ceramensis, S. tumulosa A* and *S. tumulosa B*. In our study, however, we noted that one *S. tumulosa A* (D324) grouped with *S. tumulosa B* in DAPC and it was clearly admixed in STRUCTURE analysis. This specimen was placed in sp. A in prior RAD-Seq analyses but had the same *mtMutS* and *28S* barcodes as sp. B (Quattrini et al., 2019). Perhaps these two groups of *tumulosa* have recently diverged and alleles have not yet completely sorted, and so the incongruent placement depending on marker used is not that surprising. One other incongruence with prior RAD-Seq results (Quattrini et al., 2019) was that *S. ceramensis* appeared highly admixed with *S. humilis*. This was somewhat surprising as these species were clearly distinguished by DNA barcodes and RAD-Seq data in Quattrini et al. (2019). Although it is not clear what would cause this difference, perhaps it is due to hybridization. Hybridization between these two species was not tested in Quattrini et al. (2019), but it is not impossible as hybridization is an important evolutionary mechanism in the diversification of *Sinularia*.

Species delimitation results for clade 5C were also congruent with prior RAD-Seq analyses (Quattrini et al. 2019), but there were also a few exceptions. Analyses in Quattrini et al. (2019) clearly indicated the presence of eight putative species: *S. acuta/abrupta, S. densa, S. exilis, S. lochmodes, S. maxima, S. penghuensis, S. slieringsi/penghuensis*, and *S. wanannensis*. Our results initially indicated the presence of only six putative species, grouping *S. exilis, S, maxima*, and *S. wanannensis* together, but upon further analyses at *K*=8, *S. maxima* separated from the other two species. In Quattrini et al. (2019), *S. exilis* grouped separately from the other two species in DAPC analysis, but the STRUCTURE analysis suggested that it was equally admixed between *S. densa, S. maxima*, and *S. wanannensis*. Quattrini et al. (2019) discussed that a more complete picture would emerge by including more *S. exilis* individuals and including the missing morphospecies in clade 5C. The few differences noted between this study and Quattrini et al. (2019) are more likely a reflection of the recent divergence of *Sinularia* combined with rampant hybridization and inadequate taxon sampling of the clade, rather than the method used (i.e., RAD-Seq vs UCE target-capture) to delimit species.

### Future Considerations

Our newly enhanced bait set designed to target UCEs and exons in octocorals works well to delimit among closely-related species and populations of octocorals, and because it targets some of the same loci as the hexacoral-v2 bait set (Cowman et al., 2020) it can be used to address deep to shallow level evolutionary questions across the anthozoan tree of life. However, a few methodological considerations would improve the use of this locus set in future studies. Although the mean alignment length was 1,139 ± 639 bp per locus, a few of the recovered alignments were very long (18 alignments >4,000 bp across both *Alcyonium* and *Sinularia* clades 4 and 5C). Many of these long, edge-trimmed alignments had large indels in the flanking regions, and it might be useful to remove these and/or clip alignments to approximately the same sizes to help reduce the possibility of aligning non-homologous regions. In addition, by following the phyluce tutorial for bait design, we ensured that few to no duplicate loci would be targeted, yet in analyses of specimens that were target-enriched with the new bait set and sequenced on the NovaSeq platform, phyluce randomly discarded 200-500 “duplicate” loci per specimen. These loci were removed because they either matched multiple contigs or contigs contained multiple UCE/exon loci, which perhaps is a reflection of large contig sizes obtained as compared to previous studies (e.g., Branstetter et al., 2017; Quattrini et al., 2018). Further investigation is needed to determine whether or not these “duplicates” were discarded unnecessarily. If they were not real duplicates, then keeping them could increase the number of loci retained per specimen and lead to more complete taxon occupancy matrices for analyses. In general, exploring additional bioinformatic tools to obtain and align loci might be beneficial for future studies.

The SNP calling pipeline used herein (following Zarza et al., 2016) was successful at obtaining filtered SNPs for species delimitation and population differentiation and can readily be applied to other taxonomic systems and/or locus sets (Supplemental File 2). To avoid the possibility of including linked SNPs in analysis, we filtered our SNPs to one SNP per locus or to one SNP per 1,000 bp for locus alignments ≥ 2,000 bp (∼6% of all aligned loci were > 2,000 bp). This filter can easily be set at different numbers depending on user’s goals. Another important consideration when using SNPs in analyses is having sufficient read coverage at each locus to avoid erroneous SNP calls. We called SNPs only at positions that had a read coverage of at least 10X. Although 10X might be considered low for some types of analyses, this parameter can also easily be changed by the user. Furthermore, it is possible to increase sequencing to maximize coverage, which could be important for population genomic studies that include specimens with highly degraded DNA. Nevertheless, even with only 152 filtered SNPs, the data were sufficient to differentiate populations of *A. coralloides*, highlighting the promise of this method in population genomic studies.

## Acknowledgements

NSF #1457817 provided funding to CSM. We thank the staff members of Dongsha Atoll National Park, Dongsha Atoll Research Station (DARS), and Biodiversity Research Center, Academia Sinica (BRCAS) for assistance during the field work. A. Gonzalez helped with DNA extractions and J. Hoang helped with library preparation. M. Donaldson-Matasci and E. Bush provided feedback to KE and AP on their senior theses. We thank J. McCormack and W. Tsai for use of sonicator at Occidental College. W. Tsai also provided guidance on SNP calling. B. Faircloth provided guidance on bait re-design.

## Data Accessibility

Raw Data: SRA Genbank #SAMNXXXX

Octocoral bait set, tree, alignment and SNP files: 10.6084/m9.figshare.12061038

## Author Contributions

CSM and AMQ conceived and designed the study. KE and AP developed the SNP calling pipeline and conducted phylogenetic and species delimitation analyses with guidance from AMQ and CSM. AMQ designed the baits. KE and AP wrote preliminary drafts of the manuscript. CSM and AMQ wrote the final draft of the manuscript. All authors conducted lab work and approved the final manuscript version.

## Supplementary Data

Supplementary File 1. Bait redesign protocol.

Supplementary File 2. SNP calling pipeline

Supplementary Table 1. Assembly, locus, and collection data for specimens.

### Bait design

For UCE bait redesign, we first mapped 100 bp simulated-reads from the genomes of two exemplar taxa, *Pacifigorgia irene* and *Paragorgia stephencairnsi* (unpubl. data, see Quattrini et al., 2018) to a masked *Renilla muelleri (renil*, Jiang et al., 2019) genome. Reads were mapped, with 0.05 substitution rate, using stampy v. 1 (Lunter & Goodson, 2011), resulting in 1.6 to 1.9% of reads aligning to the *renil* genome. Any alignments that included masked regions (>25%) or ambiguous bases (N or X) or were too short (<80 bp) were removed using phyluce_probe_strip_masked_loci_from_set. An SQLite table that included regions of conserved sequences shared between *renil* and each of the exemplar taxa was created using phyluce_probe_get_multi_merge_table. This table was queried using phyluce_probe_query_multi_merge_table to find conserved loci.

Phyluce_probe_get_genome_sequences_from_bed was used to extract these conserved regions from the *renil* genome. Regions were then buffered to 160 bp by including an equal amount of 5’ and 3’ flanking sequence from the *renil* genome. A temporary set of baits was designed from these loci using phyluce_probe_get_tiled_probes; two 120 bp baits were tiled over each locus and overlapped in the middle by 40 bp (3X density). This temporary set of baits was screened to remove baits with >25% masked bases or high (>70%) or low (<30%) GC content. At this stage, we concatenated temporary baits with the baits retained from the anthozoa-v1 bait set and then removed any potential duplicates (≥50% identical over ≥50% of their length) using phyluce_probe_easy_lastz and phyluce_probe_remove_duplicate_hits_from_probes_using_lastz.

This new temporary bait set was aligned back to the genomes of *P. irene, P. stephencairnsi*, and *R. muelleri* and UCE loci of *Chrysogorgia tricaulis, Cornularia pabloi*, and *Parasphaerasclera valdiviae* (see Quattrini et al., 2018) using phyluce_probe_run_multiple_lastzs_sqlite, with an identity value of 70% and a minimum coverage of 83% (default). From these alignments, baits that matched multiple loci were removed. We then extracted 180 bp of the sequences from the alignment files using phyluce_probe_slice_sequence_from_genomes. A list containing UCE loci found in at least four of the taxa was created and a new UCE bait set was designed using phyluce_probe_get_tiled_probe_from_multiple_inputs. Using this script, 120-bp baits were tiled (3X density, middle overlap) and screened for high (>70%) or low (<30%) GC content, masked bases (>25%), and duplicates as described above. Finally, the bait set was screened against the *Symbiodinium minutum* genome (Shoguchi et al., 2013) to look for any potential symbiont loci using the scripts phyluce_probe_run_multiple_lastzs_sqlite and phyluce_probe_slice_sequence_from_genomes, with a minimum coverage of 70% and minimum identity of 60%. This UCE bait set included a total of 14,179 non-duplicated baits targeting 1,707 loci.

All methods above were performed again using transcriptome data to re-design the baits targeting exons. We mapped 100 bp simulated-reads from the transcriptomes of two exemplar taxa, *Corallium rubrum* and (Pratlong et al., 2015) and *Nephthyigorgia* sp. (Zapata et al., 2015) to the *Paramuricea biscaya* transcriptome (DeLeo et al., 2018), resulting in 14 to 62% of these reads per species aligned. Following a screening for masked regions, high/low GC content, and duplicates, a temporary exon bait set was designed. At this stage, we concatenated these baits with those retained from the anthozoa-v1 bait set and then removed any potential duplicates (≥50% identical over ≥50% of their length) using phyluce_probe_easy_lastz and phyluce_probe_remove_duplicate_hits_from_probes_using_lastz.

The temporary baits were re-aligned to the transcriptomes of *C. rubrum, Eunicea flexuosa* (unpubl. data, see Quattrini et al., 2018), *Nephthyigorgia* sp., Keratoisidinae (Zapata et al., 2015), *P. biscaya*, and UCE loci of *A. digitatum, C. pabloi, C. tricaulis*, and *P. valdiviae* (see Quattrini et al., 2018) using phyluce_probe_run_multiple_lastzs_sqlite, with an identity value of 70% and a minimum coverage of 83% (default). A list containing exon loci found in at least five of the taxa was created and a new exon bait set was designed by tiling 120-bp baits (3X density, middle overlap) and screening for high (>70%) or low (<30%) GC content, masked bases (>25%), and duplicates as described above. We also screened this bait set against the *S. minutum* genome (Shoguchi et al., 2013) to look for any potential symbionts using the scripts phyluce_probe_run_multiple_lastzs_sqlite and phyluce_probe_slice_sequence_from_genomes, with a minimum coverage of 70% and minimum identity of 60%. This exon bait set included a total of 22,821 non-duplicated baits targeting 2,142 loci.

## SNP Calling Pipeline

(To be run on all or a subset of individuals that have been processed using PHYluce tutorial 1)

1. Choose reference individual by finding the individual with the most UCE/exon contigs
  a. Find individual with most contigs to use as the reference individual `less path/to/phyluce_assembly_match_contigs_to_probes.log`
2. Create fasta of UCE/exon loci from the reference individual using phyluce programs
  a. Make snp calling directory `Mkdir snp_calling`
  b. Create config named `ref.conf` file in snp calling directory that looks like: [ref]
*\*type name of reference individual here\**
  c. Run phyluce program to create list of all loci present in the reference individual phyluce_assembly_get_match_counts **\** --locus-db path/to/uce-search-results/probe.matches.sqlite **\** --taxon-list-config ref.conf **\** --taxon-group ref **\** `--output ref-ONLY.conf`
  d. Run phyluce program to create fasta file of loci present in the reference individual `phyluce_assembly_get_fastas_from_match_counts \` `--contigs /pathto/spades-assemblies/contigs \` `---locus-db /pathto/uce-search-results/probe.matches.sqlite \` `--match-count-output ref-ONLY.conf \` `--output ref-ONLY-UCE.fasta`
3. Copy, edit and run bwa-mapping
  a. Copy reads for all individuals in taxon set into folder `cd path/to/snp_calling` `mkdir reads` `cd reads` `#for each sample:` `cp -r /path/to/clean-reads-folder/sample-name ./`
  b. Download `1a_0_bwa_mapping_loop.sh` script from https://www.dropbox.com/sh/d4r8aklcvvaklxh/AADbgmTYstREvF0stHJzF6Vza?dl=0
  c. Copy bwa mapping script into snp calling directory `cd path/to/snp_calling` `cp path/to/1a_0_bwa-mapping_loop.sh bwa-mapping.sh`
  d. Make necessary edits to bwa-mapping script `nano bwa-mapping.sh`
    - in line 5, change path to `ref-ONLY-UCE.fasta`
    - in line 8, change path to reads folder
    - in line 16, change the number after `“-f”` to be one greater than the number of “/”s present in your reads path from line 8
    - in lines 20 and 21, change `“nevadae”` to `“contigs”`
    - in line 28, change path to `Mark-Duplicates.jar`
    - in line 31, change path to `BuildBamIndex.jar`
  e. Run bwa-mapping script `nohup bash bwa-mapping.sh > bwa-mapping.log`
4. Copy, modify and run indel realigner script
  a. Download `2_indelrealigner-WLET copy.sh` script from https://www.dropbox.com/sh/d4r8aklcvvaklxh/AADbgmTYstREvF0stHJzF6Vza?dl=0
  b. Copy indel realigner script into snp calling directory `cd path/to/snp_calling` `cp path/to/2_indelrealigner-WLET copy.sh indel-realigner.sh`
  c. Make necessary edits to bwa-mapping script `nano indel-realigner.sh`
    - in line 5, change path of `R=` to `ref-ONLY-UCE.fasta`
    - in line 5, change path of `O=` to `ref-ONLY-UCE.dict`
    - in line 6, change path to `ref-ONLY-UCE.fasta`
    - in line 9, change path to `ref-ONLY-UCE.fasta`
    - in line 10, change path to `*All_dedup.bam`
    - in line 18, change the number after `“-f”` to be one greater than the number of “/”s present in your reads path from line 10
  d. Run indel-realigner script `cd /path/to/snp_calling` `nohup bash indel-realigner.sh > indel-realigner.log`
5. Download, modify and run genotype recall script
  a. Download `3_genotype-recal-WLET copy.sh` script from https://www.dropbox.com/sh/d4r8aklcvvaklxh/AADbgmTYstREvF0stHJzF6Vza?dl=0
  b. Copy indel realigner script into snp calling directory `cd path/to/snp_calling` `cp path/to/3_genotype-recal-WLET copy.sh genotype-recal.sh`
  c. Make necessary edits to bwa-mapping script `nano genotype-recal.sh`

- in line 4, change path to `ref-ONLY-UCE.fasta`
- in line 7, change path to `*realigned.bam`
- in lines 15, 83, 177, 274 and 367, change the number after `“-f”` to be one greater than the number of “/”s present in your reads path from line 7
- in lines 99, 195, 290, and 383, change the path to the `recal-bams`
  d. Run genotype-recall script `cd /path/to/snp_calling` `nohup bash genotype-recal.sh > genotype-recal.log`
6. Filter SNPs using vcftools
  a. Use vcftools to filter SNP data set produced by the genotype recall script. For example, run: `vcftools --vcf genotyped_X_samples_only_PASS_snp_5th.vcf` `--min-alleles 2 --max-alleles 2 --thin 1000 --max-missing 0.75` `--max-non-ref-af 0.99 --recode --out filtered_vcf75` Which includes the parameters:
    - `--thin 1000` to ensure that no two snps are within 1000 bp of one another (a proxy filter to get 1 snp per locus)
    - `--max-non-ref 0.99` to filter out any SNPs that are the same across all samples
    - `--min-alleles 2 --max-alleles 2` to only include biallelic sites
    - `--recode` to create a new vcf of filtered SNPs
    - `--max-missing 0.75` parameter to set a threshold for taxon completeness (e.g. 0.75 only includes out SNPs for while at least 75% of individuals have an allele thus less than 25% of individuals have missing data)
    - `--out filtered_vcf75` creates a descriptive name for the output vcf More parameters can be found here: http://vcftools.sourceforge.net/man_0112a.html
7. Format filtered snps for use in DAPC and STRUCTURE analysis
  a. Use phyluce script to reformat the filtered snps `phyluce_snp_convert_vcf_to_structure --input filtered_vcf75.recode.vcf --output sin5c_input_structure.str`

---

**Table S1.**
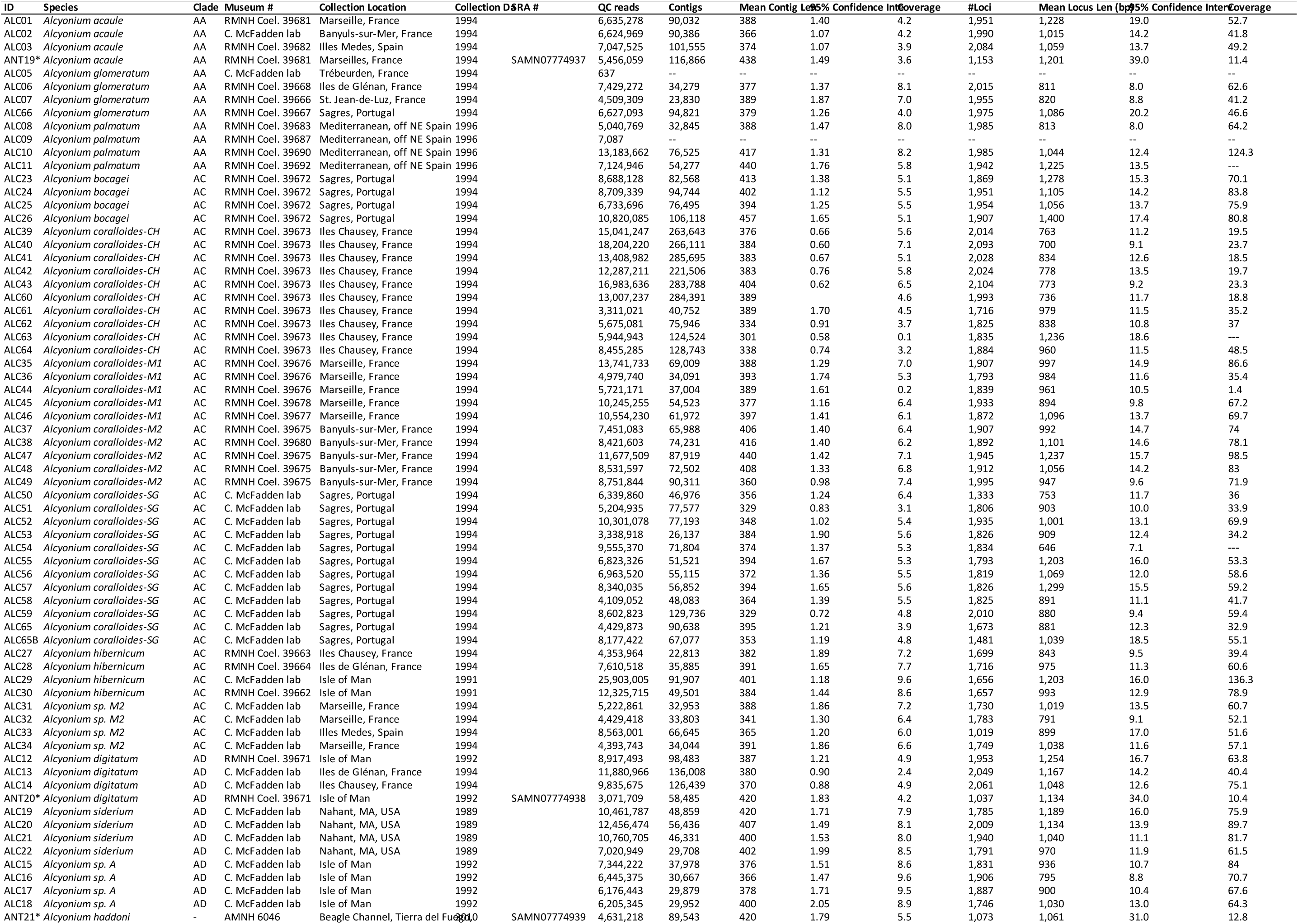
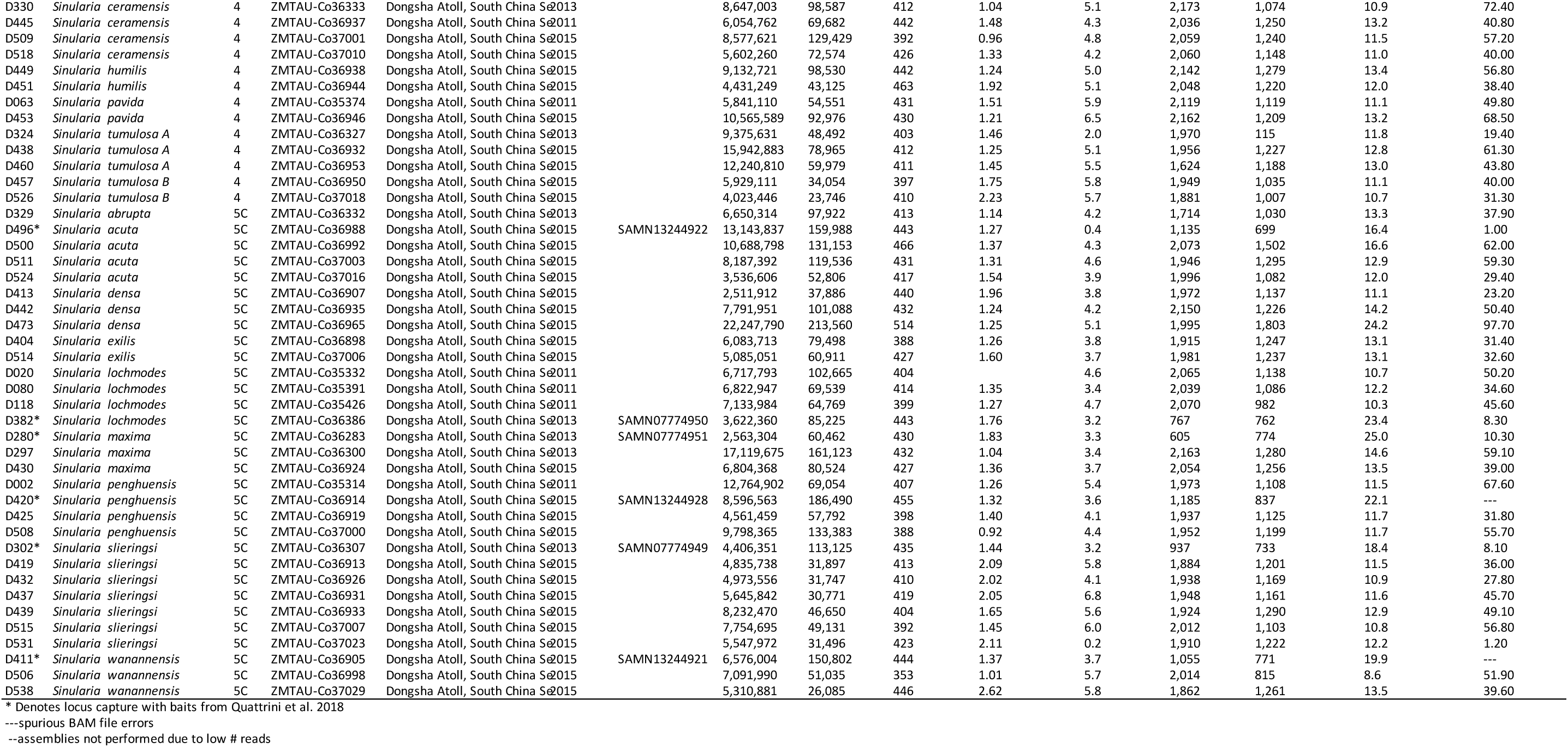
Sequencing and locus capture summary statistics.

